# Investigation of anti-IgLON5-induced neurodegenerative changes in human neurons

**DOI:** 10.1101/2020.08.27.269795

**Authors:** Matias Ryding, Mattias Gamre, Mette S Nissen, Anna Christine Nilsson, Justyna Okarmus, Anne Amalie E Poulsen, Morten Meyer, Morten Blaabjerg

## Abstract

Anti-IgLON5 disease is a progressive neurological disorder, associated with autoantibodies against a neuronal cell adhesion molecule, IgLON5. In human post-mortem brain tissue neurodegeneration and accumulation of phosphorylated-Tau is found. Whether IgLON5 antibodies induce neurodegeneration or neurodegeneration provokes an immune response remains to be elucidated. To clarify this, we exposed human stem cell derived neurons to patient anti-IgLON5 antibodies.

Human neural stem cells were differentiated for 14-48 days, and exposed from day 9-14, or day 13-48 to either 1) IgG from a patient with confirmed anti-IgLON5 antibodies; 2) IgG from a healthy person or left untreated. Electrical neuronal activity was quantified using a multi electrode array. Cultures were immunostained for β-tubulin III, phosphorylated-Tau and counterstained with DAPI. Other cultures were immunostained for synaptic proteins PSD95 and synaptophysin. Lactate dehydrogenase release and nuclei morphology analyses were performed to assess cell viability.

Spike rates and quantity of synaptic clusters were significantly reduced in anti-IgLON5 exposed cultures, and the proportion of cells with degenerative appearance was significantly higher for the anti-IgLON5 IgG group. Furthermore, the content of cells displaying accumulation of phosphorylated-Tau was higher for anti-IgLON5 exposed cultures. After five weeks, cultures exposed to anti-IgLON5 IgG contained significantly more dying cells.

In conclusion, exposure of human neurons to pathological anti-IgLON5 antibodies cause functional impairment, synaptic and neurodegenerative changes, ultimately leading to cell death. This supports autoantibodies as a causative factor responsible for the neurodegenerative changes seen in patients with anti-IgLON5 disease and mimics pathological observations.

## Introduction

Anti-IgLON5 disease is a neurological disorder first described in 2014 (Sabater *et al.*, 2014). It is thought to be an antibody-mediated disease, involving neurodegeneration (Sabater *et al.*, 2014; Gelpi *et al.*, 2016). It was initially recognized as a sleep disorder with non-rapid eye movement (NREM) and REM parasomnia, breathing dysfunction, gait instability and bulbar symptoms (Sabater *et al.*, 2014). The disease is now known to be more clinically heterogeneous in presenting symptomatology and response to immunotherapy (Gaig *et al.*, 2017; Nissen and Blaabjerg, 2019; Cabezudo-Garcia *et al.*, 2020). The disease affects men and women equally and is insidious in onset, with slow progression over several years and a fatal outcome if left untreated (Gaig *et al.*, 2017). Post-mortem analyses have revealed neuronal deposits of hyperphosphorylated Tau (p-Tau) protein in the hypothalamus and the brainstem tegmentum (Gelpi *et al.*, 2016). This deposition, consisting of three repeat and four repeat isoforms, has a different distribution compared to other neurodegenerative conditions and therefore the disorder has been described as a novel tauopathy (Braak and Braak, 1991; Braak *et al.*, 1992; Sabater *et al.*, 2014). Despite the neurodegenerative features in post-mortem brain tissue, a strong association with human leukocyte antigen haplotypes, DRB1*10:01 – DQB1*05:01, indicates an immunological association (Gaig *et al.*, 2019). The clinical manifestation of anti-IgLON5 disease was initially divided into four main phenotypes: 1) predominant sleep disorder, 2) bulbar syndrome, 3) Progressive supranuclear palsy-like syndrome and 4) cognitive impairment that may associate chorea (Gaig *et al.*, 2017). In addition, isolated gait ataxia and nervous system hyperexcitability have been included to the spectrum (Gaig and Compta, 2019). The sleep disorder is characteristic, with altered non-REM sleep initiation, vocalizations and finalistic movements (NREM parasomnia) (Sabater *et al.*, 2014; Gaig *et al.*, 2018). REM behavior disorder and sleep-breathing difficulties such as obstructive sleep apnea and stridor, can present early or late (Gaig *et al.*, 2018). Brain imaging usually shows normal or nonspecific changes on cerebral MRI with patterns of brainstem/cerebellar atrophy or hyperintensities in regions mostly affected by p-Tau (hypothalamus, brainstem, hippocampus, cerebellum) (Gaig and Compta, 2019; Nissen and Blaabjerg, 2019). CSF analysis is often normal but can show mildly increased protein levels and slight pleocytosis in the range of 5 – 10 leukocytes/μL (Nissen and Blaabjerg, 2019). (Blinder and Lewerenz, 2019). The diagnosis relies on the clinical presentation and the presence of anti-IgLON5 antibodies in serum and CSF. The precise functions of the antigen are unknown. The IgLON protein family belongs to the Immunoglobulin (Ig) protein superfamily and has three Ig-like domains, attaching to the membrane through a Glycosylphosphatidylinositol-anchor. They are expressed in synapses and engage with each other forming homo- or heterodimers (Ranaivoson *et al.*, 2019). IgLON protein dimerization involves the first Ig-domain and they seem to take part in processes such as neuronal outgrowth and synaptogenesis (Hashimoto *et al.*, 2009; Ranaivoson *et al.*, 2019). Autoantibodies directed towards IgLON5 bind specifically to an epitope in the second Ig-domain (Sabater *et al.*, 2016). This binding is thought to be pathogenic and dependent on the IgG subclass. Anti-IgLON5 IgG1 has been shown to cause irreversible internalization of IgLON5 clusters, while IgG4 may affect protein-protein interaction (Sabater *et al.*, 2016). As such, the predominant subclass may affect clinical presentation. The pathological properties of anti-IgLON5 antibodies were recently described in cultured hippocampal rat neurons with increased neurodegenerative features such as neuronal blebbing and fragmentation. However, in these cells no p-Tau deposition or cell death was observed (Landa *et al.*, 2020). To further elucidate this intriguing link between neuroinflammation and neurodegeneration, we studied the effects of patient anti-IgLON5 IgG on live human neurons derived from neural stem cells (hNSCs) and induced pluripotent stem cells (hiPSCs).

## Materials and Methods

### Ethical approval

The study was approved by the ethical committee at the Region of Southern Denmark (approval nr. S-20170134). Written informed consent was provided. All use of human stem cells was performed in accordance with the Danish national regulations, the ethical guidelines issued by the International Society for Stem Cell Research (ISSCR).

### Isolation of anti-IgLON5 IgG fractions

Serum samples containing both anti-IgLON5 IgG1 and IgG4 were obtained from a patient with a positive cell-based assay (CBA) and clinically verified anti-IgLON5 disease. The clinical features of the patient has been reported elsewhere (Nissen and Blaabjerg, 2019). A commercial CBA kit using human embryonic kidney 293 cells transfected with the IgLON5 complementary DNA and fixed with 1 % formalin (Euroimmun, Lübeck, Germany) was used to verify presence of specific IgG anti-IgLON5 antibodies. Sera were analyzed in a 1:10 dilution and titrated to endpoint (1:1,000). Presence of other encephalitis-causing autoantibodies were excluded by monospecific recombinant fixed commercial CBAs. Control human IgG was provided from sera of healthy donors and tested using commercial CBAs to exclude presence of encephalitis-causing autoantibodies.

The IgG fraction of patient and control serum were purified using a Protein A gel (MabSelect™, Amersham Biosciences) and concentrated and transferred into sterile Phosphate-Buffered Saline by centrifugation at room temperature (3000G), using protein concentrator tubes (VIVASPIN 20, Sartorius). Both purified IgG fractions from patient and controls were re-tested on CBAs and the anti-IgLON5 IgG concentration was determined (titer 1:100). The antibody eluate was stored at −80°C until usage.

### Human stem cell-derived neurons

#### hNSCs

(kindly provided by Dr. Alberto Martínez-Serrano, Autonomous University of Barcelona, Spain) were propagated and proliferated in a Poly-L-Lysine (10 ug/mL, Sigma-Aldrich) coated T-25 flask (Nunc Easyflask). Proliferation was maintained using HNSC.100 medium (composed of DMEM/F12, glucose, Hepes, AlbuMAX-I, N2 supplement, and non-essential amino acids (Gibco)), with addition of pen/strep and growth factors EGF 20 ng/ml and bFGF 20 ng/ml (R&D Systems). The medium was changed every third day. Once an adequate confluence of 80-90 % was reached, the cells were trypsinized using trypsin/EDTA and a single cell suspension was plated in 24-well plates (24 Well Cell Culture Cluster, Costar) with Poly-L-Lysine coated 15mm coverslips (Hounisen, Denmark) with a seeding density of 50.000 cells/cm^2^. Plates of monolayer cultures were incubated at 37°C: in a 5 % CO2, 95 % humidified air environment and differentiated for 14 days in HNSC.100 medium without growth factors. A 50 % medium change was conducted every third day.

#### hiPSCs

(XCell Science) predifferentiated to a neural stem cell state were propagated and proliferated in Geltrex (Gibco) coated 6-well plates. Proliferation was maintained using NSC medium composed of Neurobasal, B27, non-essential amino acids and Glutamax (Gibco) with addition of pen/strep and growth factor bFGF 10 ng/ml (R&D Systems). Medium was changed every third day and cells were enzymatically passaged using Accutase (Gibco). Cells were kept in 6-well plates until differentiation day 10 and passaged at day 5 and 10. At differentiation day 10, cells were plated in both 24-well plates with Geltrex coated 15mm coverslips and directly into 24-well plates for the multi electrode array (MEA) multi-well system.

### Exposure of human neurons to anti-IgLON5 IgG

hNSCs were divided into three treatment groups. Group I: received 2 % purified anti-IgLON5 IgG, group II: 2 % control human IgG and group III: regular media as non-exposed negative control. Cells were exposed to IgG from differentiation day 9 to 14 (Figure 1, A).

**Figure 1.**
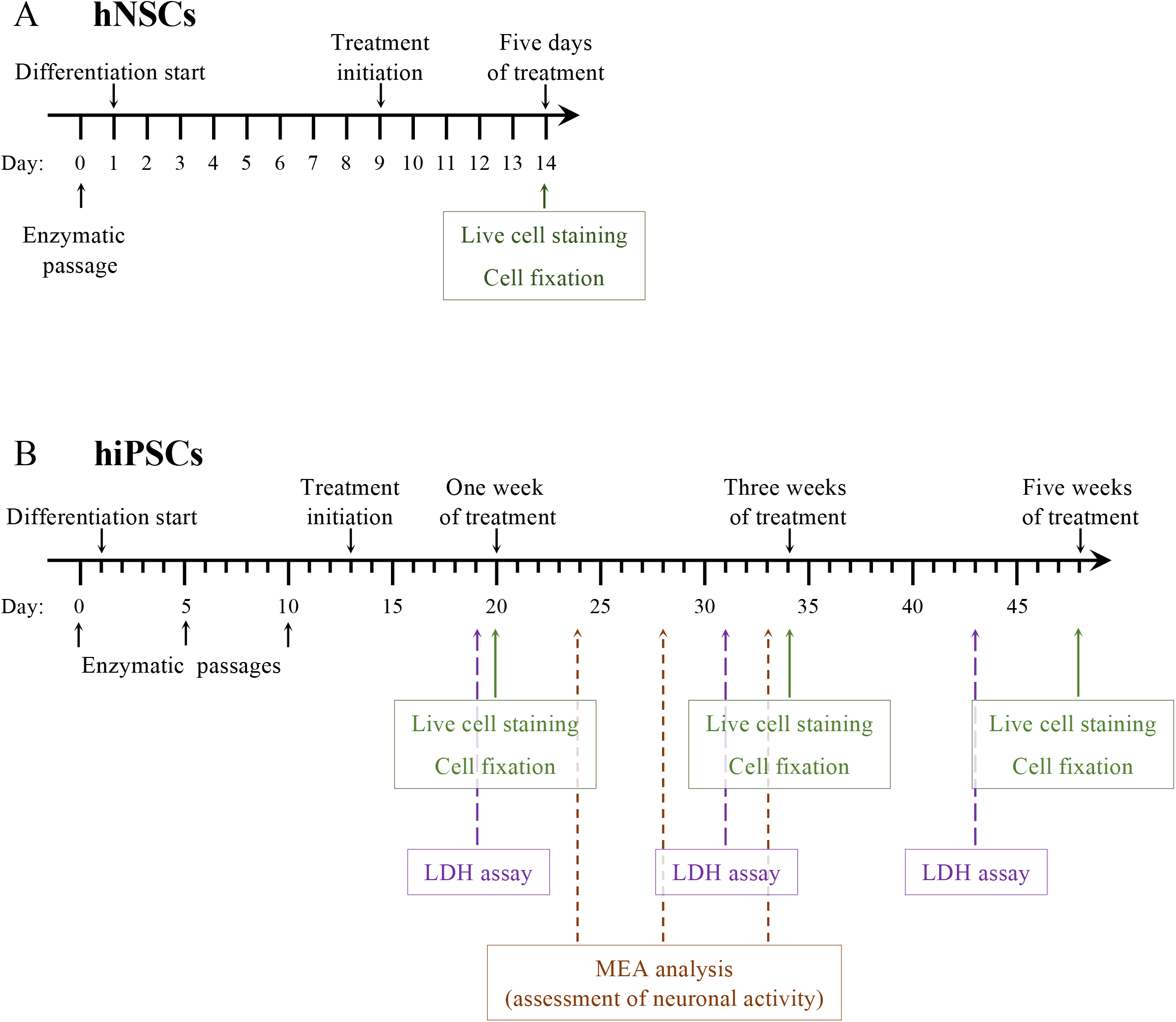
Graphical illustration of experimental design and timeline. Black arrows and text specify differentiation procedures while colored arrows and text boxes specify time points at which different analyses were performed.

hiPSCs were divided into two treatment groups: 2 % purified anti-IgLON5 IgG or 2 % control human IgG. Cells were exposed from differentiation day 13 with a treatment duration of one to five weeks. Cells to be used for immunocytochemistry were fixed in 4 % paraformaldehyde for 20 minutes and washed with Phosphate-Buffered Saline and kept at 4°C until processing. Conditioned media samples from the hiPSC cultures were collected after 6, 18 and 30 days of treatment in order to ensure equal amount of time from last media change to collection of conditioned media at each time point (Figure 1, B).

### IgLON5 live cell staining

Selected cultures were washed once in DMEM/F12 for hNSC cultures and Neurobasal Medium for hiPSC cultures followed by incubation with 2 % primary antibody (patient anti-IgLON5 IgG) in the respective culture medium for 30 min. After a second wash, cells were incubated with secondary antibody (Alexa 488, goat-anti human IgG, 1:500) for 30 min. A final washing step was performed and cultures were fixed in 4 % paraformaldehyde for 10 minutes, washed in Phosphate-Buffered Saline and mounted with ProLong Diamond Antifade (Life Technologies). Staining was performed after 2 weeks for hNSC cultures and after 1, 3 and 5 weeks of treatment for hiPSC cultures.

### Immunocytochemistry

Immunocytochemistry was performed after fixation. Cells were permeabilized with 0.1 % saponin wash buffer and blocked with a 5 % goat serum solution. Cells were then incubated with antibodies against selected proteins overnight at 4°C: Anti-β-Tubulin III (β-Tub III) (Sigma, T8660, mouse, 1:2000); anti-p-Tau (Cell Signaling, E7D3E, Rabbit, 1:100); synaptophysin (Sigma, S5768, mouse, 1:200); PSD95 (Thermo Fisher Scientific, 51-6900, rabbit, 1:1000). The following day, cells were washed and incubated with secondary antibodies dependent on the primary antibody (Alexa 555 (Thermo Fisher Scientific, goat anti-mouse 1: 500) and Alexa 488 (Thermo Fisher Scientific, goat anti-rabbit 1:500)) for one to two hrs. at room temperature (RT) followed by washing steps. A counterstain was added as 10 μM DAPI (dihydrochloride hydrate 4′,6-Diamidino-2-phenylindole) solution for 10 min. at RT. Cells were washed and mounted with ProLong Diamond Antifade.

### Image acquisition and analysis

Images were obtained using a fluorescent microscope (Olympus BX54), with a 60x objective for the live cell and synapse stainings, and a 10x objective for other immunocytochemical stainings. Five random images were collected from each 15 mm stained coverslip.

Image analysis was performed using ImageJ imaging software. For the IgLON5 live cell staining unspecific background staining was removed before the images were made binary. An automatic particle analysis gave the total number of IgLON5 clusters per image. For the total nuclear count, a Gaussian blur filter was applied before images were made binary and a watershed application segregated the individual corpuscular elements. Particle analysis gave the final nuclear count. During cell counting, a cell counter plugin tool was used to assist manual counting of pyknotic and fragmented nuclei. For analysis of neurodegeneration, neurons that contained multiple or localized swelling (blebbing) separate from a distally located growth cone, or fragmentation of the neuronal process, were counted. The percentage of neurons containing degenerative changes was calculated. In neurite outgrowth analysis, β-Tub III pictures were uploaded in NeuronJ plugin application as 8-bit images. Color balance was optimized and standardized for all images. Tracings were added for assessment of the number and length of neuronal neurites, expressed as pixel length.

For p-Tau analysis of hNSC cultures and hiPSC cultures after one week of treatment, the number of neurons containing accumulations of p-Tau protein in processes was counted with the cell counter plugin tool. Due to the complexity of the hiPSC cultures after three weeks of treatment it was not possible to do manual cell counting. Instead, the intensity of P-tau staining was measured and normalized to the intensity of the β-Tub III in the same image. The P-tau expression was then normalized to the respective control group for the hiPSC one and three weeks treatment groups for comparison of the different time points.

Quantifications of PSD95 and synaptophysin stainings were performed with the “Find Maxima” command in ImageJ.

### Neuronal spike rate analysis

Neuronal spike rate was detected using a MEA system (MultiChannel Systems). Cells were plated onto a 24 well plate with each well containing 12 electrodes. After 11, 15 and 20 days of exposure, the spike rates of the cultures were measured during a 20 minutes time period. The data was optimized by low and high frequency filters in Multiwell Analyzer (MultiChannel Systems).

### Lactate dehydrogenase (LDH) assay

An commercially available absorbance-based LDH kit (Promega) was utilized to measure the release of LDH to culture medium as an indicator of necrosis. 50 μl conditioned medium from each sample was added to wells in a 96-well plate in combination with 50μl CytoTox 96^®^ reagent followed by 30 min. at RT, and protected from light. The reaction was stopped by addition of 50μl Stop Solution to each well and absorbance was read at 490 nm to give an indirect measure of the amount of released LDH.

### Statistical analysis

For analyses of two groups at a single timepoint, data were analyzed using a two-tailed unpaired t-test. For analyses of three groups at a single timepoint (hNSC cultures) data were analyzed using a One-way, non-parametric ANOVA and *post hoc* application of Tukey’s multiple comparisons test. For analyses of two groups at two or three timepoints (hiPSC cultures), data were analyzed using a Two-way ANOVA followed by Sidak’s multiple comparisons test. Values in bar charts are presented as mean ± SEM. *P*-values below 0.05 were considered significant. All experiments were performed in independent duplicates or triplicates. Statistical analysis was performed using GraphPad PRISM, version 6.0.

### Data availability

Data supporting the findings of this study are available within the figures and the results section. Other data and extended description of methods are available from the corresponding author upon request.

## Results

### IgLON5 cluster internalization

Live cell staining for IgLON5 visualized IgLON5 clusters in the cultures (Figure 2A, B). Quantification of IgLON5 clusters in hNSC cultures revealed a significant reduction for anti-IgLON5 IgG treated cultures compared to control IgG (Control IgG: 106 ± 14 clusters per image, anti-IgLON5 IgG: 54 ± 7 clusters per image. Figure 2C)). In hiPSC cultures there was no significant difference after one week of exposure (Control IgG: 54 ± 5 clusters per image, anti-IgLON5 IgG: 35 ± 4 clusters per image, Figure 2D) but after three and five weeks of exposure, highly significant reductions of IgLON5 clusters were detected (3 weeks: Control IgG: 343 ± 40 clusters per image, anti-IgLON5 IgG: 72 ± 8 clusters per image. 5 weeks: Control IgG: 512 ± 39 clusters per image, anti-IgLON5 IgG: 169 ± 17 clusters per image).

**Figure 2.**
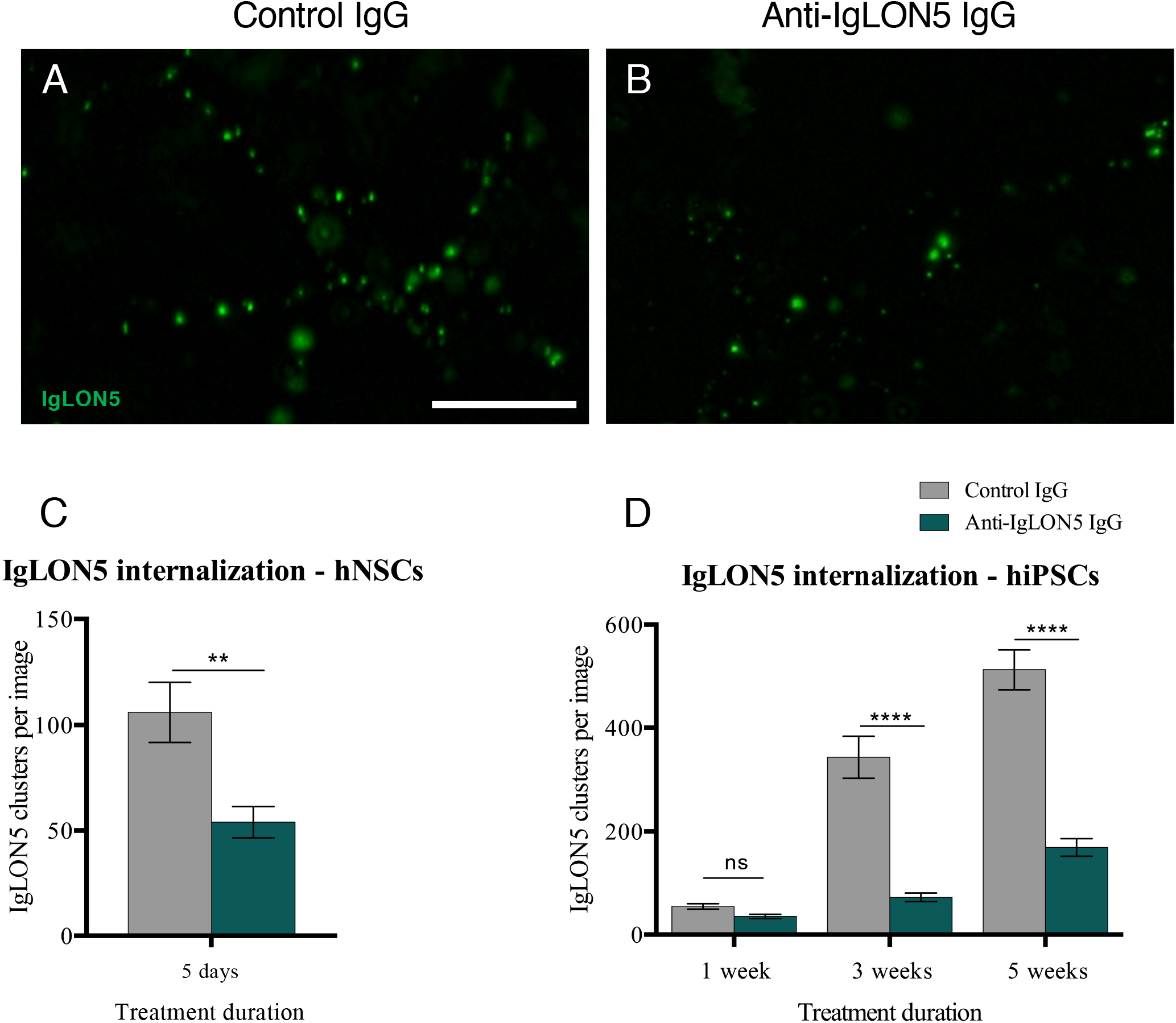
Patient antibodies reduce the content of IgLON5 clusters in human stem cell-derived neurons. Indirect immunofluorescent staining of hiPSC cultures treated with control IgG (A) or anti-IgLON5 IgG (B). Quantification of IgLON5-positive clusters in hNSC cultures showed a significant reduction of clusters when treated for 5 days with anti-IgLON5 IgG (C, *n* = 30) and a similar effect was observed in hiPSC cultures after three and five weeks of exposure (D, n = 45). Statistical analysis: two-tailed unpaired t-test (C) and Two-way ANOVA followed by Sidak’s multiple comparisons test. (D) (**: *p* < 0.01, ****: p < 0.0001). Scale bar length: 5 μm

### Multi electrode array analysis

The spike rates of hiPSC cultures were measured after 11, 15 and 20 days of antibody exposure. After 20 days, the spike rate of anti-IgLON5 IgG treated cultures was significantly reduced compared to control cultures (Control IgG: 0.720 ± 0.203Hz, Anti-IgLON5 IgG: 0.073 ± 0.032Hz, Figure 3A).

**Figure 3.**
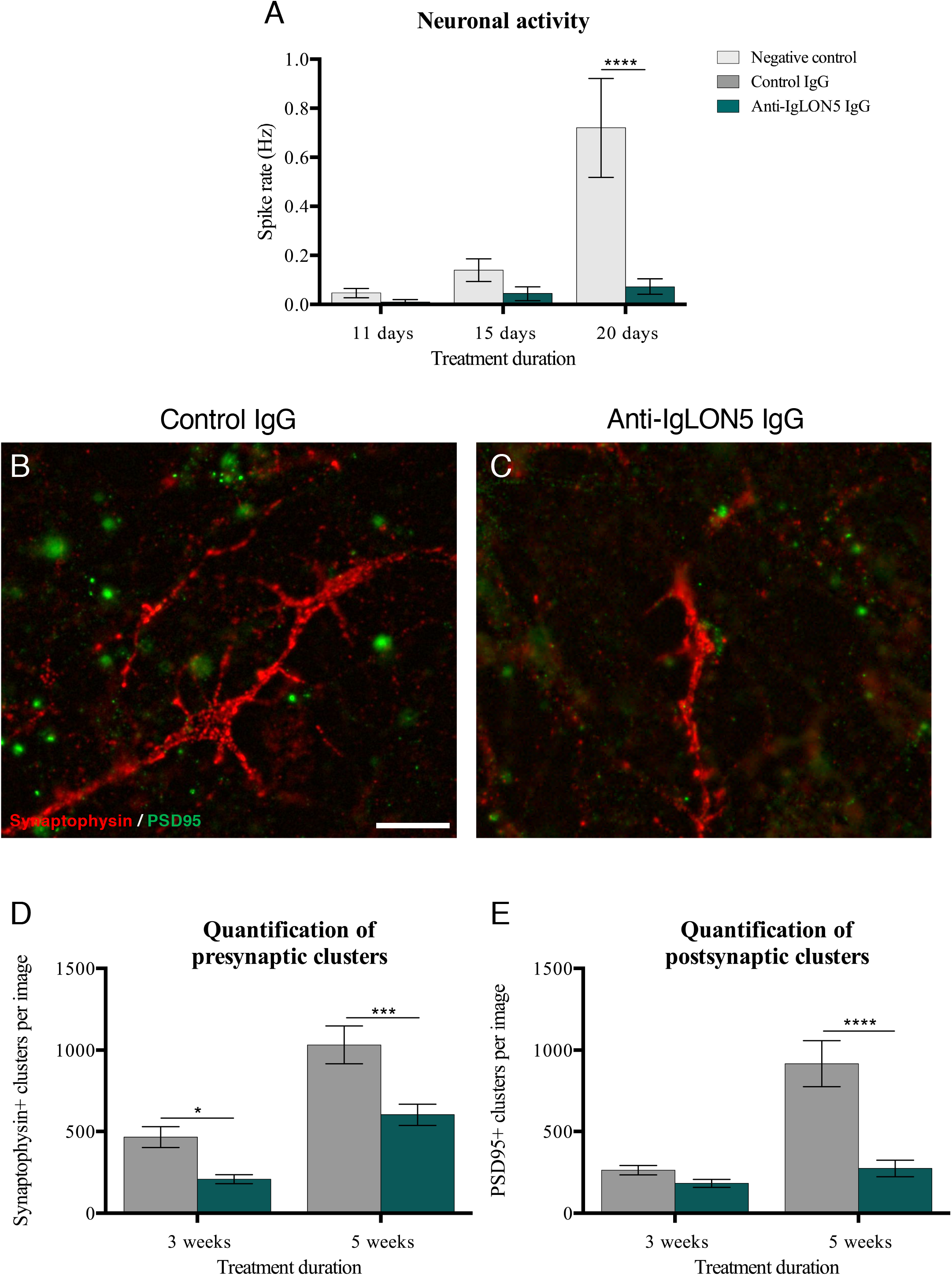
Patient antibodies reduce neuronal activity (spiking), possibly through synaptic disintegration. MEA analysis of hiPSC cultures showed the expected increase in neuronal activity as they matured, however, the spike rate of neurons treated with anti-IgLON5 IgG was significantly lower than that of untreated neurons (A, n=5). Immunostaining for the synaptic proteins synaptophysin and PSD95 (B-C) revealed a decrease in synaptophysin positive clusters after three weeks of exposure and in both synaptophysin and PSD95 clusters after five weeks of exposure (D-E, n=15). Statistical analysis: Two-way ANOVA followed by Sidak’s multiple comparisons test. (*: *p* < 0.05, ***: *p* < 0.001 ****: *p* < 0.0001). Scalebar length: 10μm

### Synapse quantification

Pre- and postsynaptic clusters were quantified in hiPSC cultures after 1, 3 and 5 weeks of treatment by immunostaining for the synaptic proteins synaptophysin (presynaptic) and PSD95 (postsynaptic) (Figure 3B, C). Analysis revealed a significant reduction of presynaptic clusters after three weeks (Control IgG: 465 ± 64 clusters per image, anti-IgLON5 IgG: 208 ± 28 clusters per image), and in both pre- and postsynaptic clusters after five weeks of antibody exposure (Control IgG: 1032 ± 116 presynaptic clusters per image and 917 ± 140 postsynaptic clusters per image, anti-IgLON5 IgG: 603 ± 66 postsynaptic clusters per image and 274 ± 50 postsynaptic clusters per image, Figure 3D, E).

### Analysis of neurite outgrowth

We analyzed neurite outgrowth in hNSC cultures exposed to anti-IglON5 IgG, control IgG or unexposed controls. No differences between exposure groups in the length or number of primary or secondary neurites or neurite endpoints were seen (data not shown).

### Early neurodegenerative changes – neuronal blebbing and fragmentation

To investigate possible early neurodegenerative changes, we obtained random images of hNSC cultures in wells immunostained with β-Tub III and analyzed the number neurons displaying axonal blebbing and/or a fragmented appearance (Figure 4A, B). The average percentage of cells with blebbing processes was significantly higher for cultures exposed to patient anti-IgLON5 IgG (14.27 ± 1.40 %), when compared to neurons exposed to control IgG (9.03 ± 0.92 %) or unexposed controls (9.26 ± 0.90 %) (Figure 4D). In addition, more fragmentation of neuronal processes was seen for IgLON5 IgG treated cultures compared to controls (IgLON5 IgG: 8.15 ± 0.71 %; Control IgG: 4.3 ± 0.46 %; Neg. Control: 4.19 ± 0.46 %) (Figure 4E).

**Figure 4.**
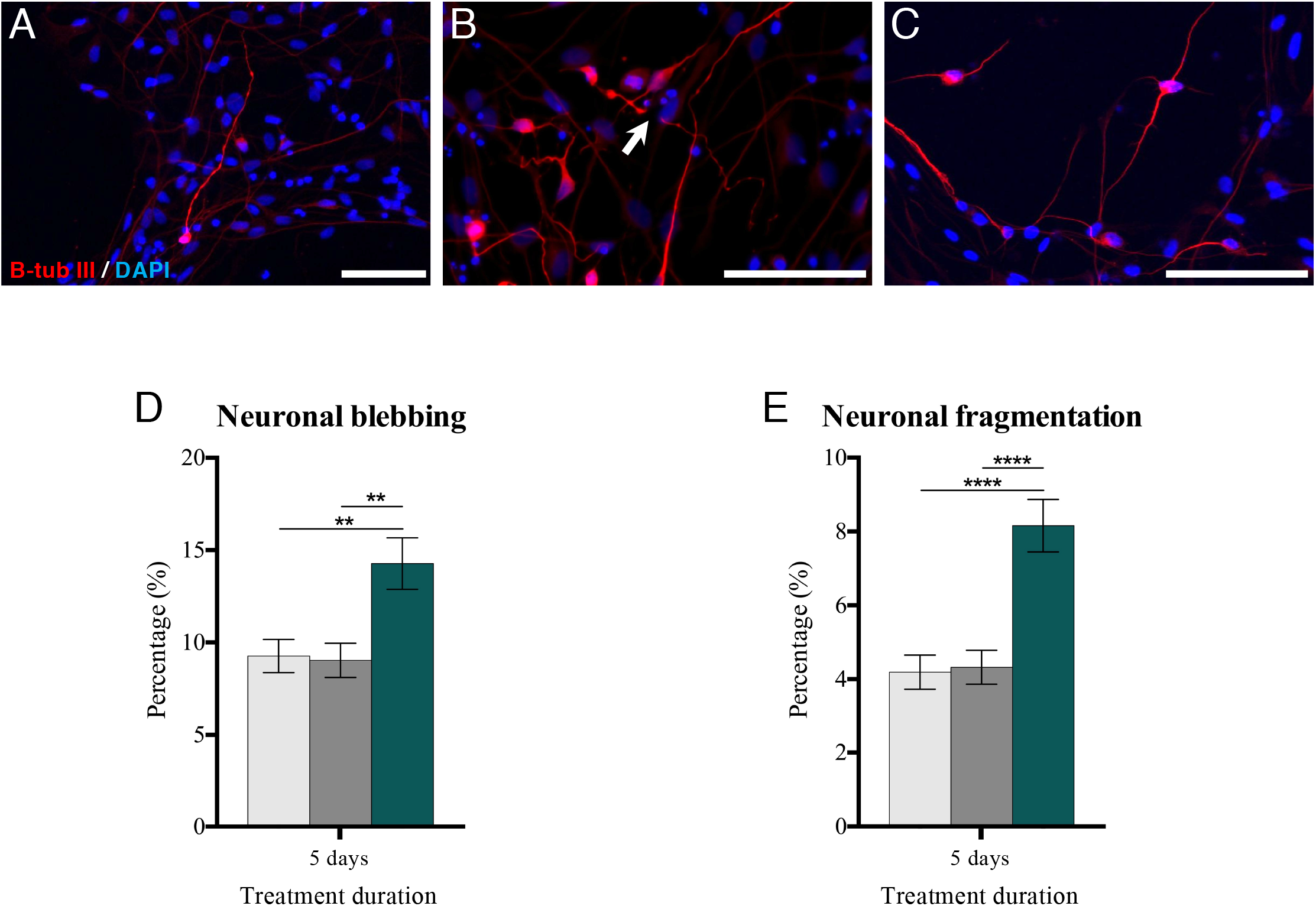
Patient antibodies cause neuronal degenerative changes. To evaluate degenerative changes in hNSC cells after 5 days of antibody exposure, cultures were immunostained with β-Tub III and analyzed. A significant difference in neuronal blebbing (A, B, D) was observed when comparing anti-IgLON5 IgG to controls. In addition, there was a significant difference in fragmented processes (B arrow, D) between anti-IgLON5 IgG, Control IgG and negative control. Representative image of neurons with normal appearance (C). Statistical analysis: One-way ANOVA followed by Tukey’s multiple comparisons test. (**: *p* < 0.01; ****: *p* < 0.0001). (Anti-IgLON5 *n* = 99; Control IgG *n* = 100; Neg. Control *n* = 94). Scale bar lengths: A, 50 μm; B-C 100 μm

### Accumulation of p-Tau protein

We continued the investigation of degenerative changes by quantifying the content of p-Tau protein in the neuronal processes of hNSC- and hiPSC-derived neurons after one and three weeks of exposure (Figure 5A, B). In hNSC cultures, we found that the relative content of neurons with p-Tau accumulation was higher for the group exposed to anti-IgLON5 IgG compared to control IgG (8.63 ± 0.30 % and 5.57 ± 0.24 %, respectively) and unexposed controls (6.07 ± 0.29 %) (Figure 5C). In hiPSC cultures the expression of p-Tau was also significantly higher in anti-IgLON5 IgG treated cultures both after one and three weeks (one week: 199 ± 14.0 % of control IgG treated cultures, three weeks: 248 ± 42.5 % of control IgG treated cultures, Figure 5D).

**Figure 5.**
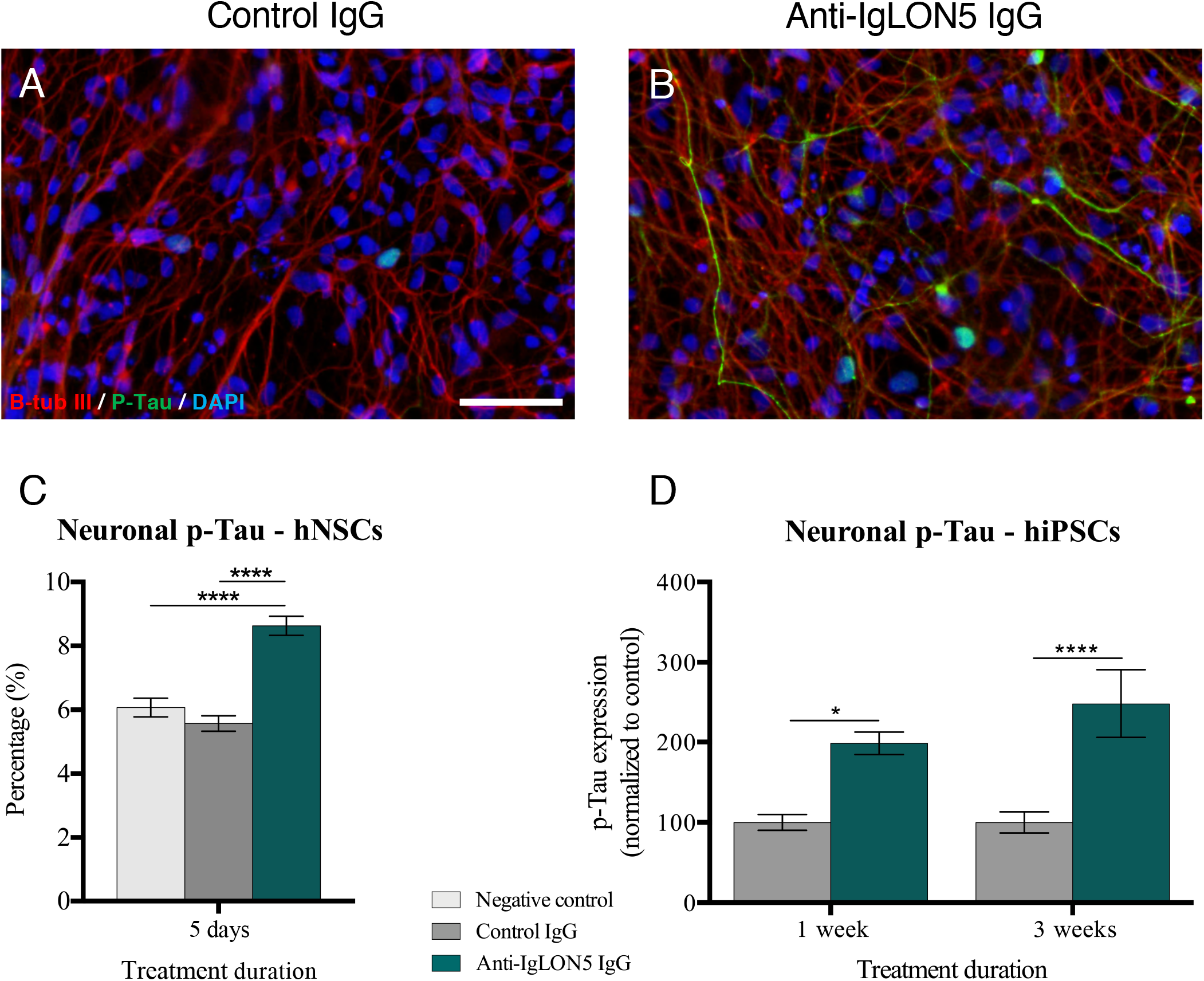
Exposure to patient antibodies cause increased accumulation of phosphorylated-Tau protein. Analysis of p-Tau in cultures. Representative images of hiPSC cultures after three weeks of exposure to control IgG (A) and anti-IgLON5 IgG (B). After 5 days exposure, hNSC cultures treated with anti-IgLON5 IgG showed a drastic increase in percentage of neurons with p-Tau when compared to cultures exposed to control IgG and negative control (C, *n* = 90-95). A similar increase in p-Tau in hiPSC cultures after one and three weeks of anti-IgLON5 IgG exposure was observed (D, n = 34-40). Statistical analysis: One-way ANOVA followed by Tukey’s multiple comparisons test (C) and Two-way ANOVA followed by Sidak’s multiple comparisons test. (*: *p* < 0.05, ****: *p* < 0.0001). Scale bar: 50 μm

### Analysis of cell viability

To investigate cell viability, we quantified nuclei with either healthy, fragmented or pyknotic morphology (Figure 6A, B, C). No differences in cell death were observed for hNSC cultures (data not shown). Similarly we found no difference in cell death for hiPSC cultures after one week of treatment. After three weeks of treatment there was a tendency towards more cells with fragmented or pyknotic morphology in anti-IgLON5 IgG treated cultures, and after five weeks, this difference was increased and statistically significant (Control IgG: 12.3 ± 1.0 % fragmented and 3.3 ± 0.3 % pyknotic nuclei, anti-IgLON5 IgG: 23.4 ± 2.3 fragmented and 5.3 ± 0.5 % pyknotic nuclei, Figure 6D, E). We further investigated necrotic cell death in hiPSC cultures by analyzing the release of LDH to culture medium as a measure of necrosis after 6, 18 and 30 days of treatment (Figure 6F). No significant differences were seen after 6 and 18 days of treatment but after 30 days of treatment, the amount of LDH in the conditioned medium from anti-IgLON5 IgG treated cultures was significantly higher than that from control IgG treated cultures, (Control IgG: 0.75 ± 0.02A, anti-IgLON5 IgG: 0.86 ± 0.04A).

**Figure 6.**
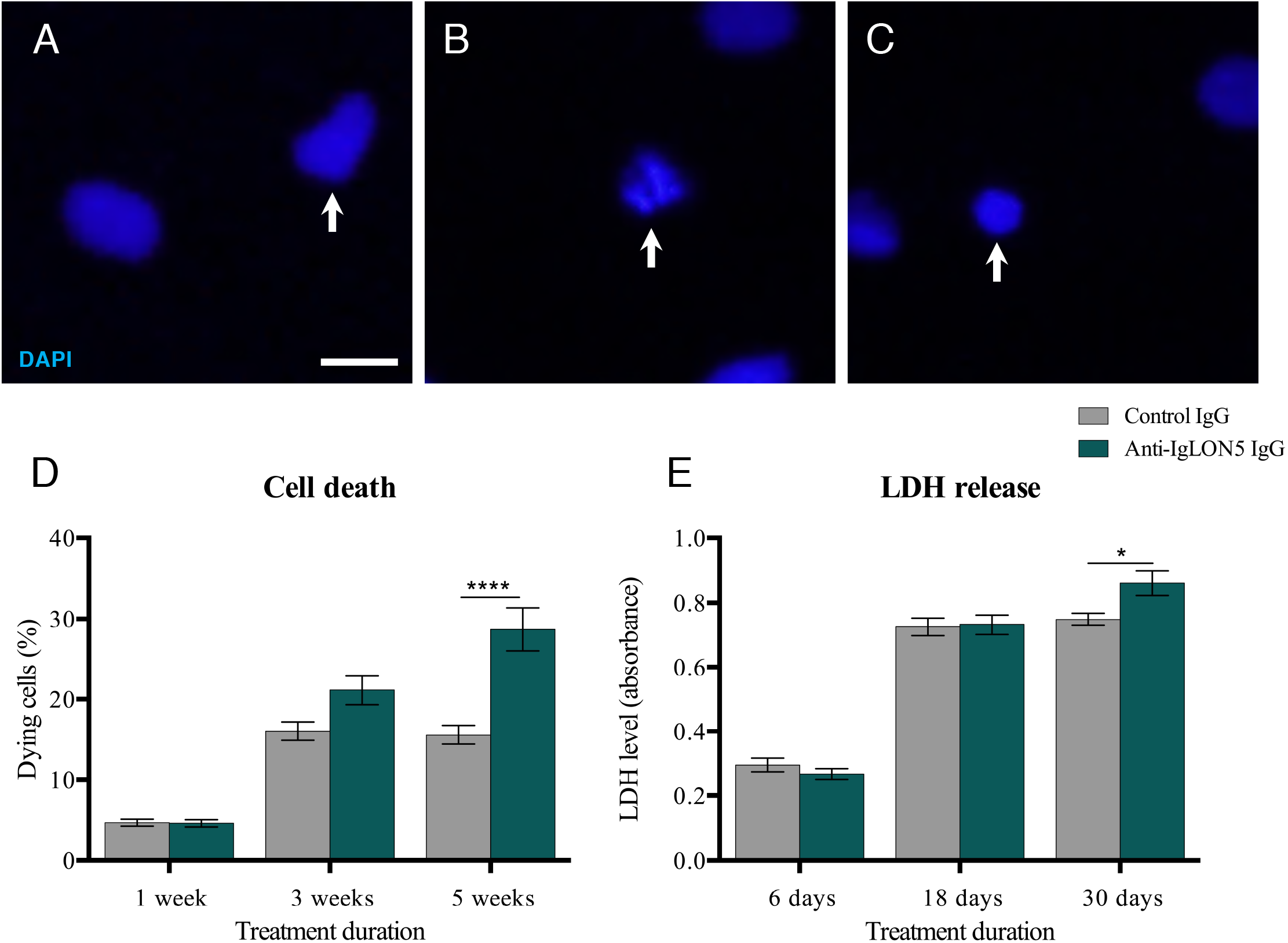
Patient antibodies caused increase cell death in hiPSC-derived neural cultures. Examples of cell nuclei with healthy (A), fragmented, (B) or pyknotic (C) morphology in hiPSC cultures. Quantification of nuclei with unhealthy morphology revealed increased content of dying cells after five weeks of anti-IgLON5 IgG exposure. A LDH-assay (a marker of necrosis) showed increased LDH content in conditioned medium from anti-IgLON5 IgG treated cultures after five weeks of exposure (E). Statistical analysis: Two-way ANOVA followed by Sidak’s multiple comparisons test. (*: *p* < 0.05, ****: *p* < 0.0001). Scale bar: 10 μm

## Discussion

In this study, we describe the neuropathological effects induced by anti-IgLON5 antibodies providing the first demonstration of these effects on living human neurons.

In both hNSC and hiPSC cultures, anti-IgLON5 IgG reduced the amount of extracellular IgLON5 clusters. This is in accordance with previous reports on mechanism of patient anti-IgLON5 antibodies (Sabater *et al.*, 2016; Landa *et al.*, 2020). Patient antibodies also reduced the electrophysiological activity of neurons and although this has never before been shown as an effect of anti-IgLON5 antibodies, changes in electrophysiological properties is a commonly described effect of antibodies of other subtypes of AE (Dalmau *et al.*, 2008; Peng *et al.*, 2015; Petit-Pedrol *et al.*, 2018). However, unlike most targets of AE antibodies, the synaptic function of IgLON5 is unknown. As described above, the IgLON protein family are intercellular adhesion molecules with a recognized role in synaptogenesis and neuronal outgrowth, but very little is known about the role of IgLON5 (Hashimoto *et al.*, 2009; Ranaivoson *et al.*, 2019). To further investigate this, we quantified clusters of post- and presynaptic proteins (PSD95 and synaptophysin, respectively) and found them both to be significantly reduced in cultures exposed to anti-IgLON5 IgG. This indicates that IgLON5 is indeed involved in synaptogenesis or synaptic stability, which may be the mechanistic explanation of the observed decreased neuronal communication.

Neuronal cells with an unhealthy morphology were recognized in all hNSC cultures. We found that cultures exposed to anti-IgLON5 IgG displayed a significantly higher percentage of neurons with degenerative changes such as axonal blebbing and fragmentation. This is in line with recent observations in cultured rat hippocampal neurons and further provides evidence that exposure to anti-IgLON5 antibodies causes neurodegeneration (Landa *et al.*, 2020). Moreover, these cells were found to have smaller/pyknotic nuclei and were seen through a spectrum, where they eventually released their neuronal processes. It is tempting to speculate that these cells were destined to die and if given more time would add to the total and specific neuronal cell death in the anti-IgLON5 exposed cell cultures. Axonal blebbing has been used to describe an unhealthy neuronal morphology in primary hippocampal and cortical rat cultures (Loehfelm *et al.*, 2020) and as a marker for chronic degenerative changes caused by neuroinflammation in *in vivo* murine models (Chakravarty *et al.*, 2020).

A neurite outgrowth assay is commonly used to assess neuronal health and the neurotoxicity of substances at concentrations that may not affect cellular viability (Krug *et al.*, 2013). Further, it is used to investigate the growth of neurites in models of neurodegenerative diseases (Bogetofte *et al.*, 2019). Despite an unhealthy morphology, we did not find any differences in neurite number, length and branchpoints in hNSC cultures treated with anti-IgLON5 IgG compared to controls. Whether this is due to a very short antibody exposure period or because IgLON5 does not regulate neurite outgrowth remains unclear, as it was impossible to assess in hiPSC cultures after extended antibody exposure due to the complexity of the cultures.

A characteristic feature of neurodegeneration is the accumulation of p-Tau protein in aggregates (Saha and Sen, 2019). Patients with anti-IgLON5 disease are found to have accumulation of p-Tau protein in post-mortem examinations (Gelpi *et al.*, 2016). Whether anti-IgLON5 antibodies induce this neurodegenerative feature or these neurodegenerative changes provoke an immune response, leading to secondary antibody formation, is still debated. In hNSC cultures the content of neuronal p-Tau was significantly higher when exposed to anti-IgLON5 IgG. This finding supports the hypothesis that neuroinflammation precedes neurodegeneration and that anti-IgLON5 antibodies contribute to the neurodegenerative changes found in patients with anti-IgLON5 disease. We found no increase in cell death in hNSC cultures treated with anti-IgLON5 IgG, which may be due to the short duration of exposure. In hiPSC cultures P-tau accumulation was present early after exposure (1 week) and was further increased after three weeks. Furthermore, five weeks of exposure to anti-IgLON5 IgG induced an increase in cell death as seen by the elevated fraction of fragmented and pyknotic nuclei and the increased LDH release. To our knowledge, this is the first demonstration of antibody mediated degenerative changes in cultures of human neurons and a strong indicator that the autoantibodies in anti-IgLON5 encephalitis are causative for the disease and not a secondary event.

Anti-IgLON5 disease is rare with an estimated incidence of 1/150 000 and with only two cases documented in Denmark (Nissen and Blaabjerg, 2019). One Danish patient has died and thus we were limited to use serum samples from only one single patient. Our study is further limited by the use of serum samples with a combination of mainly IgG4 and less IgG1. However, it reflects the combined effect of antibodies in our patient and further studies of monoclonal antibody subtypes could help to dissect the cellular mechanisms leading to accumulation of p-Tau and morphological changes, seen in our neural cultures.

This study raises several important implications demonstrated for the first time: 1) patient anti-IgLON5 antibodies caused a decrease in synaptic proteins and consequently in neuronal activity; 2) human neurons exposed to patient anti-IgLON5 IgG exhibited significantly increased neuronal degenerative changes in the form of axonal blebbing, fragmentation, and accumulation of p-Tau; 3) and ultimately exposure to anti-IgLON5 antibodies lead to increased cell death (summarized in Figure 7). These findings support the hypothesis that anti-IgLON5 antibodies lead to neurodegeneration and correlate with the neuropathological findings in patients postmortem.

**Figure 7.**
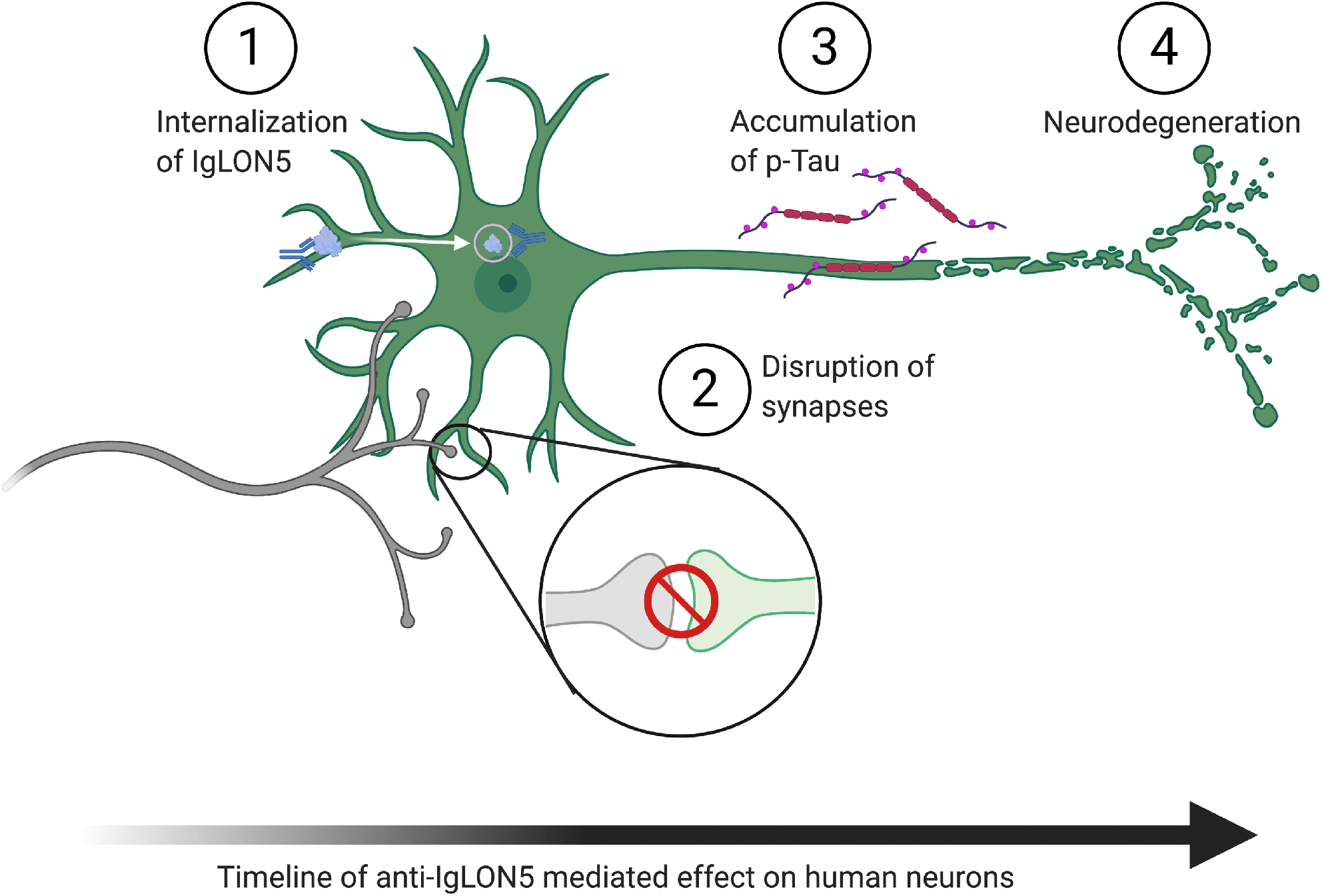
Graphical illustration of the pathological consequences of exposure to anti-IgLON5 antibodies over time. Based on our temporal data we hypothesize that after the initial IgLON5 internalization (1), synapses are disrupted (2) leading to decreased neuronal communication. This is following by accumulation of p-Tau (3), fragmentation of neurons, and subsequently neuronal cell death (4). Created with BioRender.com

## Abbreviations

CBA: Cell-based assay
hiPSC: Human induced pluripotent stem cell
hNSC: Human Neural Stem cell
Ig: Immunoglobulin
LDH: Lactate dehydrogenase
MEA: Multi-electrode array
p-Tau: Phosphorylated-Tau
RT: Room temperature

## References

Blinder T, Lewerenz J. Cerebrospinal Fluid Findings in Patients With Autoimmune Encephalitis-A Systematic Analysis. Front Neurol 2019; 10: 804.

Bogetofte H, Jensen P, Okarmus J, Schmidt SI, Agger M, Ryding M, et al. Perturbations in RhoA signalling cause altered migration and impaired neuritogenesis in human iPSC-derived neural cells with PARK2 mutation. Neurobiology of Disease 2019; 132.

Braak H, Braak E. Neuropathological stageing of Alzheimer-related changes. Acta Neuropathol 1991; 82(4): 239–59.

Braak H, Jellinger K, Braak E, Bohl J. Allocortical neurofibrillary changes in progressive supranuclear palsy. Acta Neuropathol 1992; 84(5): 478–83.

Cabezudo-Garcia P, Mena-Vazquez N, Estivill Torrus G, Serrano-Castro P. Response to immunotherapy in anti-IgLON5 disease: A systematic review. Acta Neurol Scand 2020; 141(4): 263–70.

Chakravarty D, Saadi F, Kundu S, Bose A, Khan R, Dine K, et al. CD4 Deficiency Causes Poliomyelitis and Axonal Blebbing in Murine Coronavirus-Induced Neuroinflammation. J Virol 2020; 94(14).

Dalmau J, Gleichman AJ, Hughes EG, Rossi JE, Peng X, Lai M, et al. Anti-NMDA-receptor encephalitis: case series and analysis of the effects of antibodies. Lancet Neurol 2008; 7(12): 1091–8.

Gaig C, Compta Y. Neurological profiles beyond the sleep disorder in patients with anti-IgLON5 disease. Curr Opin Neurol 2019; 32(3): 493–9.

Gaig C, Ercilla G, Daura X, Ezquerra M, Fernandez-Santiago R, Palou E, et al. HLA and microtubule-associated protein tau H1 haplotype associations in anti-IgLON5 disease. Neurol Neuroimmunol Neuroinflamm 2019; 6(6).

Gaig C, Graus F, Compta Y, Hogl B, Bataller L, Bruggemann N, et al. Clinical manifestations of the anti-IgLON5 disease. Neurology 2017; 88(18): 1736–43.

Gaig C, Iranzo A, Santamaria J, Graus F. The Sleep Disorder in Anti-lgLON5 Disease. Curr Neurol Neurosci 2018; 18(7).

Gelpi E, Hoftberger R, Graus F, Ling H, Holton JL, Dawson T, et al. Neuropathological criteria of anti-IgLON5-related tauopathy. Acta Neuropathol 2016; 132(4): 531–43.

Hashimoto T, Maekawa S, Miyata S. IgLON cell adhesion molecules regulate synaptogenesis in hippocampal neurons. Cell Biochem Funct 2009; 27(7): 496–8.

Krug AK, Balmer NV, Matt F, Schonenberger F, Merhof D, Leist M. Evaluation of a human neurite growth assay as specific screen for developmental neurotoxicants. Arch Toxicol 2013; 87(12): 2215–31.

Landa J, Gaig C, Plaguma J, Saiz A, Antonell A, Sanchez-Valle R, et al. Effects of IgLON5 Antibodies on Neuronal Cytoskeleton: A Link between Autoimmunity and Neurodegeneration. Ann Neurol 2020.

Loehfelm A, Elder MK, Boucsein A, Jones PP, Williams JM, Tups A. Docosahexaenoic acid prevents palmitate-induced insulin-dependent impairments of neuronal health. FASEB J 2020; 34(3): 4635–52.

Nissen MS, Blaabjerg M. Anti-IgLON5 Disease: A Case With 11-Year Clinical Course and Review of the Literature. Front Neurol 2019; 10: 1056.

Peng XY, Hughes EG, Moscato EH, Parsons TD, Dalmau J, Balice-Gordon RJ. Cellular Plasticity Induced by Anti-alpha-Amino-3-Hydroxy-5-Methyl-4-Isoxazolepropionic Acid (AMPA) Receptor Encephalitis Antibodies. Annals of Neurology 2015; 77(3): 381–98.

Petit-Pedrol M, Sell J, Planaguma J, Mannara F, Radosevic M, Haselmann H, et al. LG antibodies alter K-v I.I and AMPA receptors changing synaptic excitability, plasticity and memory. Brain 2018; 141: 3144–59.

Ranaivoson FM, Turk LS, Ozgul S, Kakehi S, von Daake S, Lopez N, et al. A Proteomic Screen of Neuronal Cell-Surface Molecules Reveals IgLONs as Structurally Conserved Interaction Modules at the Synapse. Structure 2019; 27(6): 893–906 e9.

Sabater L, Gaig C, Gelpi E, Bataller L, Lewerenz J, Torres-Vega E, et al. A novel non-rapid-eye movement and rapid-eye-movement parasomnia with sleep breathing disorder associated with antibodies to IgLON5: a case series, characterisation of the antigen, and post-mortem study. Lancet Neurol 2014; 13(6): 575–86.

Sabater L, Planaguma J, Dalmau J, Graus F. Cellular investigations with human antibodies associated with the anti-IgLON5 syndrome. J Neuroinflamm 2016; 13.

Saha P, Sen N. Tauopathy: A common mechanism for neurodegeneration and brain aging. Mech Ageing Dev 2019; 178: 72–9.

